# SuperSurv: A Unified Framework for Machine Learning Ensembles in Survival Analysis

**DOI:** 10.64898/2026.03.11.711010

**Authors:** Yue Lyu, Steven Hsesheng Lin, Xuelin Huang, Ziyi Li

## Abstract

This paper introduces **SuperSurv**, a user-friendly R package for building, evaluating, and interpreting ensemble models for right-censored survival data. Although many survival modeling methods are available, existing tools are often model-specific and lack a unified platform for systematically integrating, comparing, and ensembling heterogeneous learners. **SuperSurv** addresses this gap by providing a unified interface for diverse survival learners, including models that return full survival curves as well as methods that produce only risk scores. All learner outputs are mapped to calibrated survival probability curves on a common evaluation time grid, enabling direct comparison and ensemble construction across heterogeneous model classes.

**SuperSurv** implements stacking of survival models using inverse-probability-of-censoring weighted (IPCW) Brier risk to estimate ensemble weights in the presence of right censoring. The framework integrates hyperparameter tuning, time-dependent benchmarking metrics, and visualization tools for survival model evaluation. In addition, the package provides post-hoc interpretability utilities based on SHAP(SHapley Additive exPlanations) values and supports covariate-adjusted restricted mean survival time (RMST) contrasts through g-computation.

By bridging the gap between theoretical rigor and clinical application, **SuperSurv** offers researchers a comprehensive ecosystem for modern survival analysis. The **SuperSurv** package is open-source and available on CRAN at https://CRAN.R-project.org/package=SuperSurv and on GitHub at https://github.com/yuelyu21/SuperSurv. An empirical example using the METABRIC breast cancer dataset demonstrates a complete workflow from model training and benchmarking to explainability and clinically interpretable survival contrasts.

## 1. Introduction

Supervised learning for time-to-event outcomes is a cornerstone of precision medicine and public health, where accurate predictions drive individualized risk assessment. Traditionally, the Cox proportional hazards (PH) model (Cox 1972) has been the dominant analytical tool. However, its reliance on strict log-linear and proportional hazards assumptions often renders it inadequate for capturing the complex data structures present in modern clinical datasets (Therneau and Grambsch 2000).

To address these limitations, the statistical learning community has developed a vast array of survival-adapted algorithms. These span from non-parametric tree-based methods—such as Random Survival Forests (Ishwaran, Kogalur, Blackstone, and Lauer 2008; Wright and Ziegler 2017) and Accelerated Oblique Random Forests (Jaeger, Welden, Lenoir, Speiser, Segar, Pandey, and Pajewski 2024)—to powerful gradient and component-wise boosting machines (Chen and Guestrin 2016; Binder and Schumacher 2008). Furthermore, the landscape has expanded to include regularized regressions (Simon, Friedman, Hastie, and Tibshirani 2011), flexible generalized additive models (Wood 2017), survival support vector machines (Fouodo, König, Weihs, Ziegler, and Wright 2018), and Bayesian additive regression trees (Sparapani, Logan, McCulloch, and Laud 2016).

Despite this proliferation of methods, it is rarely known *a priori* which specific algorithm or hyperparameter configuration will perform best for a given dataset. Rather than committing to a single parametric or algorithmic specification, predictive accuracy can be systematically optimized by combining candidate models using an ensembling framework.

### 1.1. The Super Learner and the Survival Challenge

The Super Learner (SL) provides an asymptotically optimal stacking framework for selecting a best-performing learner or an optimally weighted ensemble from a prespecified library by minimizing an out-of-sample loss function via *V* -fold cross-validation (Van der Laan, Polley, and Hubbard 2007; Polley, Rose, and van der Laan 2011). Originally developed for fully observed continuous or binary outcomes (e.g., via the **SuperLearner** package), extending SL to right-censored time-to-event data presents the following major methodological and computational challenges.

First, because outcomes are right-censored, traditional loss functions (such as standard mean squared error) cannot be directly applied. Performance objectives and cross-validation procedures must incorporate Inverse Probability of Censoring Weighting (IPCW) (Graf, Schmoor, Sauerbrei, and Schumacher 1999; Gerds and Schumacher 2006) to remain consistent. Over the past decade, the statistical literature has steadily evolved to address this challenge:

- **Discrete-Time Stacking:** Early theoretical work by Polley and van der Laan (2011) extended SL by reframing continuous survival time as a sequence of discrete-time hazard classification tasks. While innovative, this approach requires artificially expanding the dataset across time intervals, which becomes computationally prohibitive for high-dimensional clinical data.
- **Restricted Continuous-Time Stacking:** Golmakani and Polley (2020) subsequently proposed a continuous-time Super Learner; however, they restricted the candidate library strictly to proportional hazards-based algorithms, such as basic Cox regression, Elastic Net, and Lasso-Cox. This formulation structurally excluded modern, non-parametric machine learning engines.
- **Generalized Continuous-Time Stacking:** More recent breakthroughs generalized the continuous-time framework to accommodate arbitrary machine learning algorithms. Westling, Luedtke, Gilbert, and Carone (2024) established a robust cross-fitted IPCW objective for drawing inference from treatment-specific survival curves using diverse learners.
- **Joint Ensembling:** Building on the need for robust censoring weights, Munch and Gerds (2024) introduced the “joint survival super learner,” demonstrating the theoretical necessity of simultaneously optimizing both the event-time algorithms and the censoring-distribution algorithms.

Second, even with robust continuous-time IPCW loss functions in place, survival algorithms fundamentally differ in their *prediction targets*. Some methods—such as classical parametric models, Kaplan-Meier estimators, and Random Survival Forests (Ishwaran *et al*. 2008)— natively output an estimated survival curve 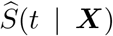 evaluated across a predefined time grid. Conversely, many of the highly predictive modern machine learning engines—such as Gradient Boosting Machine (e.g., **xgboost**), Support Vector Machine (e.g., **survivalsvm**), and penalized regression (e.g., **glmnet**)—are optimized via partial likelihood and output only a relative risk score or linear predictor *η*(***X***) without a fully specified baseline hazard. This architectural mismatch creates a critical implementation gap. Without an additional, model-agnostic mathematical calibration step, risk-score learners cannot be directly stacked alongside curve-based learners. Furthermore, naïve combinations of predictions originating from such disparate algorithmic sources are not guaranteed to be properly scaled as probabilities, directly comparable, or even monotonically decreasing over time *t*.

Third, even if a diverse survival ensemble is successfully constructed, its clinical application is often hindered by a lack of interpretability and difficulties in evaluating treatment efficacy. High-dimensional ML ensembles are inherently “black boxes,” making it difficult for clinicians to interpret individual patient risk factors. Furthermore, extracting a population-level treatment effect from a highly non-linear ensemble is challenging. While classical analysis relies on the Hazard Ratio (HR), the HR is non-collapsible and often misleading when the true underlying data exhibits crossing hazards over time, which can be naturally captured by ML algorithms (Martinussen and Vansteelandt 2013). Therefore, researchers may prefer absolute, collapsible measures such as the restricted mean survival time (RMST). By applying G-computation (standardization) to the ensemble survival predictions (Uno, Claggett, Tian *et al*. 2014), **SuperSurv** supports covariate-adjusted marginal contrasts on the RMST scale. When the binary grouping variable corresponds to a manipulable intervention and standard identification assumptions hold, the resulting standardized contrast may also admit a causal interpretation; for non-manipulable attributes such as sex or biomarker status, it should instead be interpreted as an adjusted marginal contrast.

Despite the rapid development of machine learning methods for survival analysis, the practical software ecosystem remains fragmented. Individual packages typically implement a single model class or a narrow family of algorithms, making it difficult to combine heterogeneous learners within a unified ensemble framework. In practice, analysts must manually reconcile incompatible prediction formats, time grids, and risk-score outputs before ensemble learning can be applied.

### 1.2. The Fragmented Software Landscape

While the theoretical foundations of both discrete-time and continuous-time survival Super Learning have been established, the current R (R Core Team 2024) software ecosystem for survival ensemble learning remains fragmented. Translating these methodological advances into accessible, unified software has proven challenging, leaving applied researchers with several distinct but incomplete paradigms:

1. **Object-Oriented ML Frameworks:** Packages like mlr3proba (Sonabend, Király, Bender, Bischl, and Lang 2021) provide an extensive machine learning ecosystem for survival analysis. However, it relies heavily on R6 object-oriented syntax, presenting a steep learning curve. Furthermore, to ensemble modern risk-score-based learners (like xgboost) alongside curve-based learners, users must manually construct complex reduction pipelines (e.g., `distrcompositor`) to estimate the baseline hazard; it does not happen automatically.
2. **Classification-Based Stacking:** Recent frameworks like survML (Wolock, Gilbert, Simon, and Carone 2024) avoid IPCW entirely by recasting survival estimation as a pooled binary classification problem (“global survival stacking”) evaluated over a discrete time grid. While this allows the use of standard binary classifiers, it artificially expands the dataset size across the time grid (incurring heavy computational costs) and precludes the direct ensembling of native, specialized survival algorithms (e.g., Random Survival Forests).
3. **Parametric-Focused Ensembles:** The survivalSL package (Sabathe and Foucher 2024) recently introduced a practical survival Super Learner. However, its utility is constrained by a library focused predominantly on classical parametric models (e.g., AFT models with Weibull or Gamma distributions) and standard Cox penalizations. Because it lacks an automated calibration engine, it cannot seamlessly integrate the raw risk scores of modern “black-box” ML engines.
4. **Methodological Codebases:** The foundational continuous-time SL papers are accompanied by research codebases, such as the survSuperLearner GitHub package (Westling *et al*. 2024) and the jossl scripts (Munch and Gerds 2024). While mathematically rigorous, they function primarily as proof-of-concept repositories rather than production-ready software. Both are fundamentally constrained by extremely limited built-in algorithm libraries. For instance, survSuperLearner supports a library of models that natively output survival curves (e.g., Cox, Weibull, Random Survival Forests) but structurally excludes modern risk-score-based algorithms (e.g., gradient boosting, SVM) because it lacks an automated baseline hazard calibration blueprint. Similarly, jossl provides native support for only three algorithms. Furthermore, neither ecosystem provides native tools for Explainable AI (XAI) or the estimation of covariate-adjusted marginal treatment effects.

These limitations motivate the development of SuperSurv, an R package designed to unify survival ensemble learning, model calibration, and interpretation within a single extensible framework. Recent tutorial(Keogh, Diaz-Ordaz, van Geloven, Gran, and Tanner 2025) and applied work(Tanner, Sharples, Daniel, and Keogh 2021; Li, Chien, Lyu *et al*. 2025) suggests that Super Learner methods for time-to-event outcomes are becoming increasingly relevant in real-world survival analyses; however, existing implementations often remain bespoke, requiring analysts to manually construct wrappers and limiting the breadth of learner libraries that can be combined in practice.

### 1.3. The SuperSurv Package: Design and Contributions

The **SuperSurv** package is designed to eliminate this software fragmentation. Building conceptually on the continuous-time framework of Westling *et al*. (2024), **SuperSurv** provides the robust software infrastructure necessary to transform methodological theory into a comprehensive, user-facing predictive ecosystem.

Crucially, **SuperSurv** extends beyond pure prediction to directly address the interpretability and treatment evaluation gaps identified above. By integrating Explainable AI (XAI) directly into its architecture(Lundberg and Lee 2017), the package bridges complex ensembles to the **fastshap** (Štrumbelj and Kononenko 2014; Greenwell 2024) and **survex** (Spytek, Krzyzński, Langbein, Baniecki, Wright, and Biecek 2023) ecosystems. Additionally, the package provides native G-computation routines to estimate covariate-adjusted Average Treatment Effects (ATE) via the Restricted Mean Survival Time (RMST) (Uno *et al*. 2014), entirely bypassing the non-collapsible Hazard Ratio.

By bridging the gap between theoretical rigor and precision medicine application, **SuperSurv** offers the following core components:

- **Model-Agnostic Calibration:** a plug-in procedure that maps arbitrary risk scores *η*(***X***) to calibrated survival curves via Breslow-type baseline hazard recovery, enabling machine learning models that produce only risk scores to be integrated seamlessly with models that estimate full survival functions. This allows modern ML (e.g., **xgboost, survivalsvm**) to be ensembled seamlessly alongside classical models.
- **Dual-Objective Stacking:** Convex weight estimation by minimizing user-specified IPCW Brier risk or IPCW Negative Log-Likelihood (Log Loss), a feature absent in foundational survival SL packages.
- **Unified API and Expanded Library:** A standardized API representing predictions as absolute survival curves on a user-defined time grid, harmonizing 19 distinct base algorithms and 6 automated high-dimensional screening algorithms.
- **Covariate-Adjusted RMST Contrasts: SuperSurv** provides post hoc estimation of standardized group contrasts on the restricted mean survival time (RMST) scale, yielding clinically interpretable summaries on an absolute time scale.
- **Interpretability Pipelines: SuperSurv** provides native support for global and local model interpretation, including SHAP-based explanations and seamless bridging to time-dependent interpretability tools through the **survex** ecosystem.
- **High-dimensional screening:** Automated variable screening algorithms (e.g., Elastic Net, Marginal Cox, Variance) to handle genomic-scale datasets.
- **Evaluation Utilities:** A built-in visualization and benchmarking suite for time-dependent Brier scores(Brier 1950), Uno’s C-index(Uno, Cai, Pencina, D’Agostino, and Wei 2011), and time-dependent AUC curve(Heagerty, Lumley, and Pepe 2000; Blanche, Dartigues, and Jacqmin-Gadda 2013).

### 1.4. Organization of the paper

The remainder of this paper is organized as follows. Section 2.1 introduces the statistical notation for right-censored survival data and inverse probability of censoring weighting (IPCW). Section 2.2 describes the harmonization blueprint used to convert arbitrary risk scores into calibrated survival probabilities. Sections 2.3 and 2.4 detail the theoretical and computational representations of the dual-objective IPCW loss functions used to optimize the ensemble weights. Section 2.5 highlights how this architecture provides dynamic robustness to proportional hazards misspecification, followed by an outline of the iterative survival–censoring optimization algorithm in Section 2.6.

Shifting from methodology to the software ecosystem, Section 3 outlines the **SuperSurv** R package design, demonstrating its unified API and Explainable AI (XAI) integration. Finally, Section 4 illustrates an end-to-end empirical application of the package using the METABRIC breast cancer dataset (Curtis, Shah, Chin, Turashvili, Rueda, Dunning, Speed, Lynch, Samarajiwa, Yuan *et al*. 2012; Kvamme, Borgan, and Scheel 2019).

## 2. Statistical Framework and Ensemble Learning Architecture

This section introduces the statistical framework underlying the **SuperSurv** package, including the representation of right-censored survival data, transformation of heterogeneous model outputs to survival probabilities, IPCW loss functions for ensemble learning, and the iterative procedure used to jointly estimate survival and censoring models.

### 2.1. Right-Censored Survival Data and IPCW Notation

Let *T* denote the true event time, *C* the right-censoring time, and Δ = 𝕀 (*T* ≤ *C*) the event indicator. Let ***X*** ∈ ℝ^p^ represent a *p*-dimensional vector of baseline covariates. We observe *n* independent and identically distributed realizations of the right-censored data structure

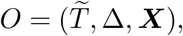

where 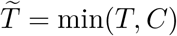 is the observed follow-up time.

We assume conditionally independent censoring,

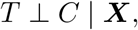

which ensures identifiability of the conditional survival distribution. Our primary predictive target is the conditional survival function

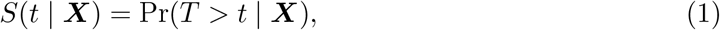

**Figure 1:**
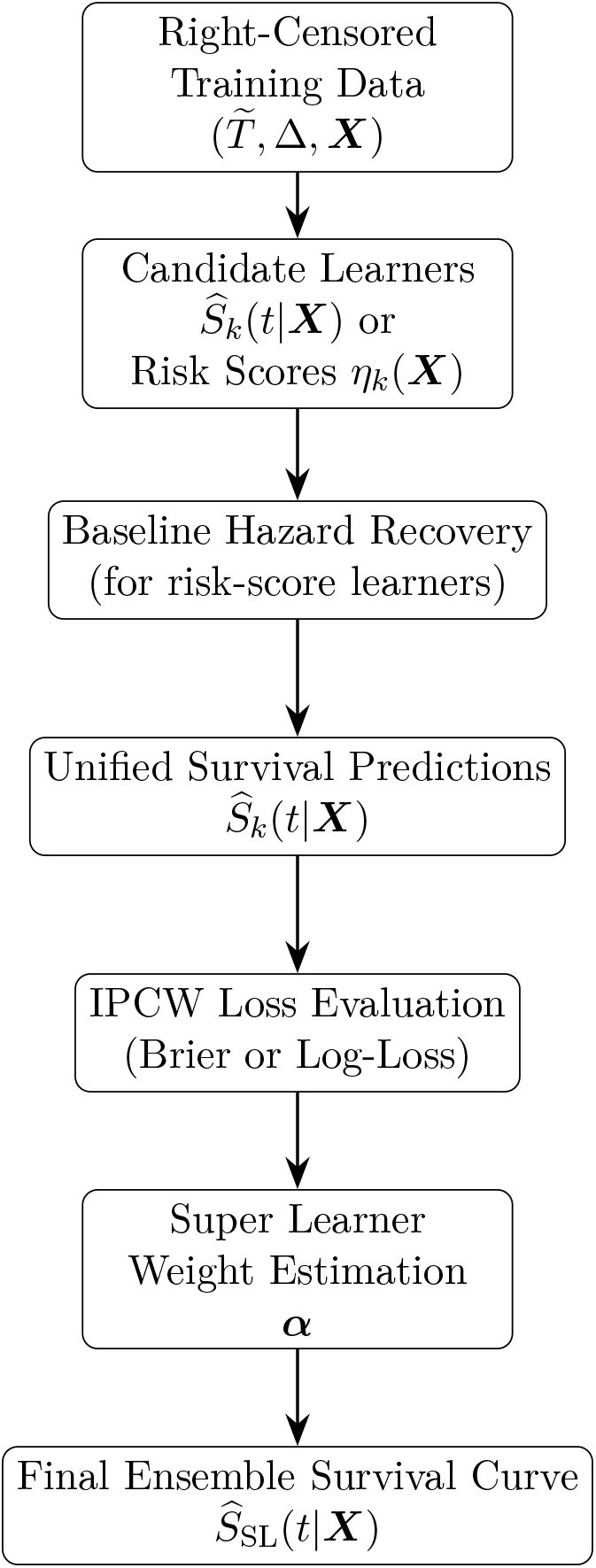
Overview of the **SuperSurv** framework. Heterogeneous survival learners are first converted to survival probability curves (if necessary), then combined using IPCW-based meta-learning to produce the final ensemble survival estimator.

which is related to the cumulative hazard function

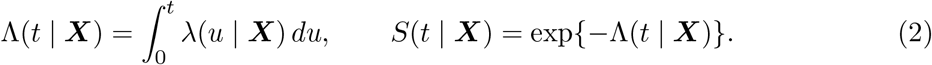

Because the event time may be censored, empirical risk functions based on fully observed outcomes cannot be applied directly. To address this issue, we employ inverse probability of censoring weighting (IPCW). Let

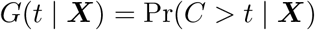

denote the conditional survival function of the censoring distribution. For a prediction evaluated at time *t*, the IPCW weight is defined as

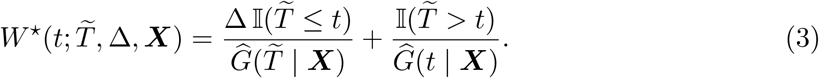

The first term corresponds to individuals who experience the event prior to time *t*, while the second term accounts for individuals who remain at risk beyond time *t*. These weights correct for informative loss of follow-up and allow censored observations to contribute appropriately to risk estimation. These weights enable unbiased estimation of prediction risk under right-censoring and form the basis of the ensemble loss functions used in Section 2.3.

In the **SuperSurv** package, this mathematical formulation maps directly to the core function interface for fitting the ensemble model.

### 2.2. Model Output Harmonization

To construct a valid Super Learner ensemble, all candidate algorithms must estimate the same target parameter,

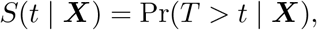

on a common evaluation time grid

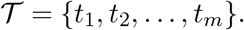

However, modern survival learning algorithms produce heterogeneous outputs. Some algorithms directly estimate the survival function, while others optimize likelihoods that yield only relative risk scores or linear predictors. Consequently, the predicted quantities must be transformed into a common representation before stacking can occur.

Let 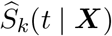 denote the survival estimate produced by learner *k*. In **SuperSurv**, every learner is wrapped into a standardized interface that returns a matrix

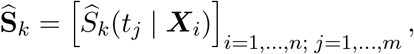

where rows correspond to individuals and columns correspond to time points in 𝒯 .

To achieve this harmonization, **SuperSurv** distinguishes between three classes of learners:

1. **Direct survival learners** that natively output survival curves.
2. **Hazard-based learners** that produce linear predictors or log-risk scores.
3. **Utility-score learners** that produce ranking scores not directly linked to the hazard scale.

Table 1 summarizes the output format of each learner currently implemented in **SuperSurv**.

**Table 1:** Output format of base learners implemented in **SuperSurv**.

| Learner | Output type | Conversion to survival curve |
| --- | --- | --- |
| Kaplan–Meier | survival probability | direct |
| Parametric AFT (Weibull, Lognormal, etc.) | survival probability | direct |
| Random Survival Forest (**ranger**, **randomForestSRC**) | survival probability | direct |
| Oblique Random Survival Forest (**aorsf**) | survival probability | direct |
| Bayesian Additive Regression Trees (**BART**) | survival probability | direct |
| Generalized Additive Model (**mgcv**) | survival probability | direct |
| Cox PH (**survival**) | log-risk | Breslow hazard recovery |
| Penalized Cox (**glmnet**, ridge) | linear predictor | Breslow hazard recovery |
| Gradient Boosting (**gbm**) | log-risk | Breslow hazard recovery |
| XGBoost Cox | log-risk | Breslow hazard recovery |
| CoxBoost | log-risk | Breslow hazard recovery |
| Survival Tree (**rpart**) | event-rate proxy | Breslow hazard recovery |
| Survival SVM | utility score | Cox calibration |

#### Baseline hazard recovery

Many machine learning survival algorithms optimize the Cox partial likelihood and therefore output only a linear predictor

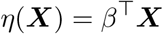

or an equivalent log-risk score. Because the partial likelihood leaves the baseline hazard *H*_0_(*t*) unspecified, such models do not directly provide absolute survival probabilities.

To recover survival probabilities, **SuperSurv** estimates the baseline cumulative hazard using a Breslow-type estimator. Let (*t*_i_, Δ_i_) denote the observed follow-up time and event indicator for subject *i*, and let

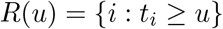

denote the risk set at time *u*.

For a learner producing linear predictors *η*_i_, the cumulative baseline hazard is estimated as

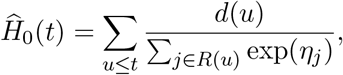

where *d*(*u*) denotes the number of events observed at time *u*.

Algorithm 1 describes the computational procedure used within **SuperSurv** to recover the baseline hazard from predicted risk scores.

#### Cox calibration for utility-score learners

Certain algorithms, such as Survival Support Vector Machines, produce utility scores that are not guaranteed to lie on the proportional hazards scale. For these learners, **SuperSurv** fits a univariate Cox proportional hazards model

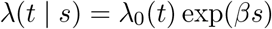

##### Algorithm 1

Baseline hazard recovery (Breslow-type updates)

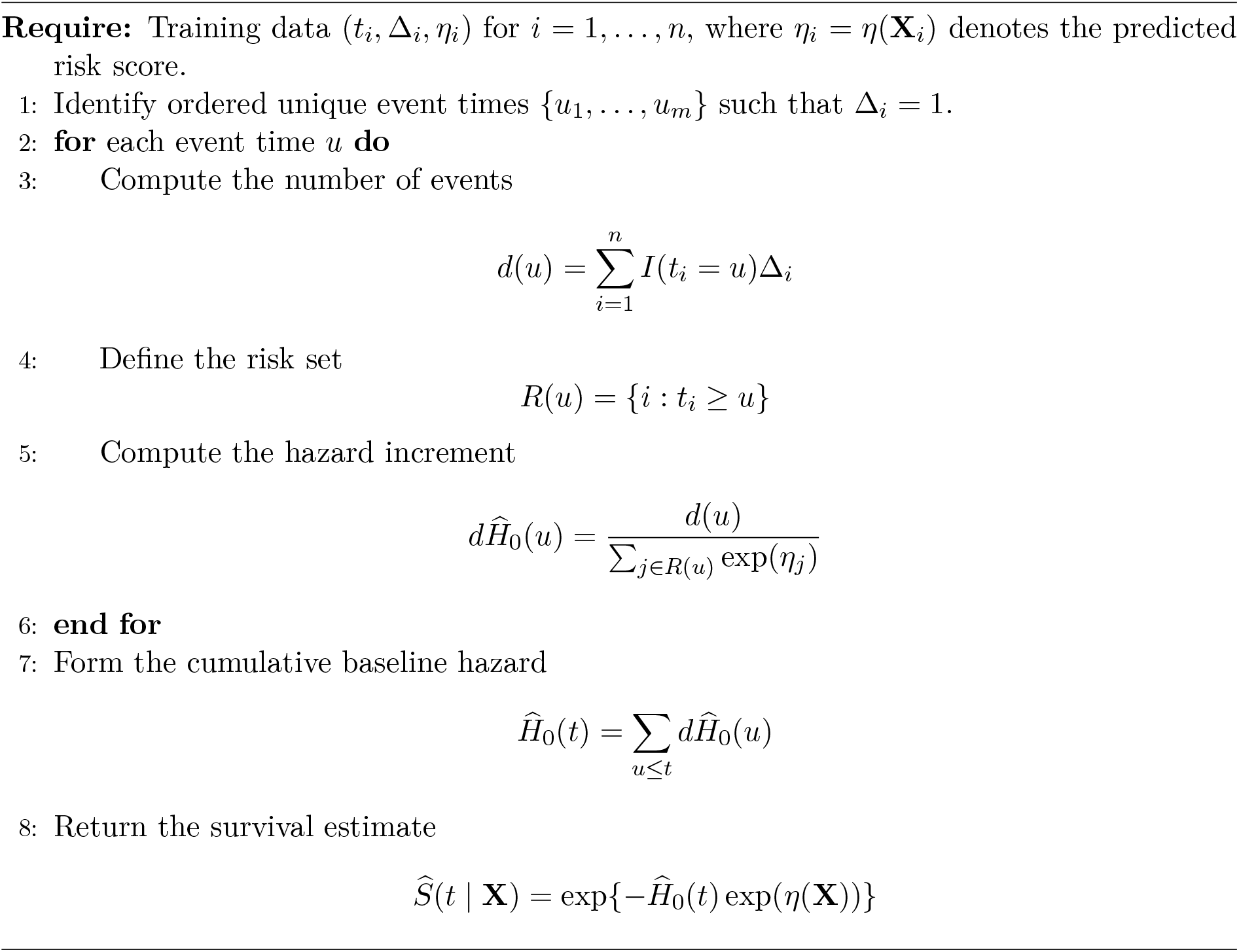

where *s* denotes the raw model score. This step calibrates the score onto the hazard scale before applying the Breslow estimator.

#### Interpolation onto a common time grid

Different learners evaluate survival curves at different time points. To ensure compatibility across models, predicted survival curves are interpolated onto the shared grid 𝒯 using a step-function interpolation

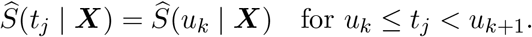

This constant interpolation preserves the monotonicity of survival curves while ensuring all learners produce predictions on the same grid.

#### Numerical stability safeguards

Several engineering safeguards are implemented to ensure numerical stability during hazard recovery:

- Linear predictors are truncated to avoid numerical overflow when computing exp(*η*).
- Survival curves are constrained to remain monotone decreasing via cumulative minima operations.
- Design matrices for training and prediction are aligned by padding missing columns to handle differing factor levels across cross-validation folds.
- Extremely small time values are truncated to prevent undefined logarithms in parametric models.

These safeguards ensure robust prediction across heterogeneous machine learning libraries. Once predictions from heterogeneous learners are expressed as survival probability curves *S*(*t* | ***X***), they can be compared and combined using a common prediction loss. The SuperSurv framework estimates ensemble weights by minimizing an inverse-probability-of-censoring weighted loss defined over these survival predictions.

### 2.3. Dual-Objective IPCW Loss Functions

The goal of the meta-learning stage is to estimate the optimal convex combination of *K* candidate learners. Let the stacked predictor be defined as

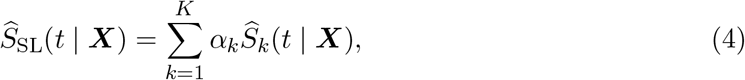

where the weight vector ***α*** = (*α*_1_, …, *α*_K_) lies on the simplex

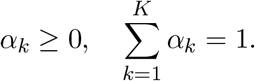

Weights are estimated via *V* -fold cross-validation by minimizing an inverse-probability-of-censoring weighted (IPCW) prediction loss.

#### IPCW Brier-Loss

The default objective minimizes the IPCW brier-loss, which forms the basis of the Super Learner for conditional survival distributions developed by Westling *et al*. (2024). For a prediction *S*(*t* | ***X***) at time *t*, the loss is defined as

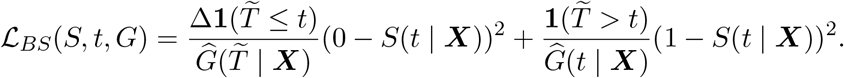

This formulation corresponds to an inverse-probability-of-censoring weighted squared error between the predicted survival probability and the true survival status at time *t*.

While the integrated Brier score provides the theoretical foundation for survival distribution stacking (Westling *et al*. 2024), modern machine learning systems often rely on log-loss objectives due to their strong probabilistic interpretation and sensitivity to miscalibrated predictions. In the binary outcome setting, log-loss is equivalent to the cross-entropy loss widely used in machine learning. To accommodate this perspective, **SuperSurv** additionally implements an IPCW log-loss objective for estimating ensemble weights.

Compared with squared-error loss, log-loss penalizes overconfident but incorrect survival predictions more strongly, which may improve ensemble calibration in practice.

#### IPCW Log-Loss

In addition to the integrated Brier score objective, **SuperSurv** provides an alternative meta-learning criterion based on IPCW log-loss. This loss emphasizes probabilistic accuracy by penalizing confident but incorrect predictions more strongly, making it particularly suitable when well-calibrated survival probabilities are required. The IPCW log-loss is defined as

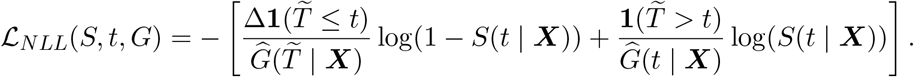

The ensemble weights are obtained by minimizing the empirical IPCW log-loss

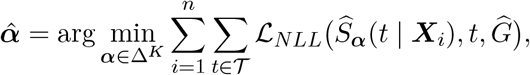

where Δ^K^denotes the simplex of nonnegative weights that sum to one.

To our knowledge, this represents one of the first practical implementations of an IPCW cross-entropy objective for survival distribution stacking within a Super Learner framework.

While the loss functions above are defined for a fixed time point, the **SuperSurv** framework evaluates prediction error across an entire time horizon. In practice, the continuous-time loss is approximated using a discrete evaluation grid, which leads to the computational representation described below.

### 2.4. Computational Representation of the Loss Function

Following Westling *et al*. (2024), the Super Learner for survival outcomes is defined through the minimization of an expected loss for the conditional survival distribution. For the weighted integrated Brier score, the population loss is defined as

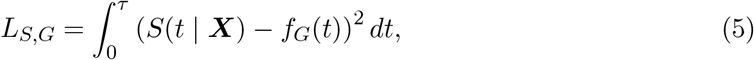

where

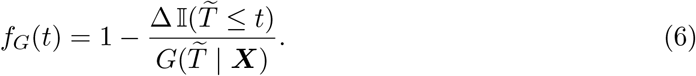

Here 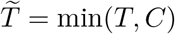 denotes the observed follow-up time, Δ = 𝕀 (*T* ≤ *C*) is the event indicator, and *G*(*t* ***X***) denotes the conditional censoring survival function. The function *f*_G_(*t*) serves as an inverse-probability-of-censoring weighted pseudo-outcome that corrects for right censoring. In practice, the integral over time cannot be evaluated analytically and must be approximated numerically. **SuperSurv** therefore evaluates predictions on a predefined time grid

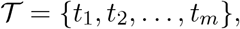

and approximates the continuous-time loss using a discrete summation over this grid.

Let 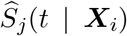 denote the predicted survival probability for observation *i* at time *t* from candidate learner *j*. The empirical loss used for model comparison is then computed as

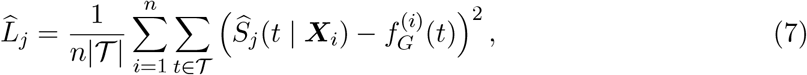

where 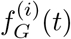 denotes the IPCW pseudo-outcome for observation *i* evaluated at time *t*. This formulation corresponds directly to the matrix-based implementation used in the package, where survival predictions are represented as an *n ×* |*𝒯* | matrix and the loss is evaluated element-wise across observations and time points.

The asymptotic properties of the Super Learner estimator under this loss function, including oracle inequalities and consistency results, are established in Westling *et al*. (2024). The implementation in **SuperSurv** follows this theoretical framework while providing a computationally efficient discrete-time approximation suitable for large-scale machine learning ensembles.

### 2.5. Robustness to Proportional Hazards Misspecification

A key advantage of the Super Learner framework is its ability to combine learners that rely on different structural assumptions about the hazard function.

Many classical survival models, including Cox proportional hazards models and their penalized extensions, assume that the hazard function factorizes as

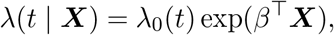

which implies proportional hazards across covariate strata. While this assumption simplifies estimation and interpretation, it can be violated in complex biomedical datasets where treatment effects or covariate influences vary over time.

The **SuperSurv** library intentionally includes both proportional hazards models (e.g., **coxph, glmnet, xgboost** Cox objectives) and flexible machine learning estimators that do not rely on proportional hazards assumptions (e.g., Random Survival Forests, Bayesian Additive Regression Trees, and survival trees).

Because ensemble weights are estimated by minimizing cross-validated prediction risk, the Super Learner automatically adapts to the underlying hazard structure. When the proportional hazards assumption holds, Cox-type learners typically achieve low prediction error and receive higher ensemble weights. Conversely, when proportional hazards are violated, flexible nonparametric learners tend to achieve better predictive performance, causing the ensemble weights assigned to Cox-type models to shrink toward zero.

This adaptive weighting mechanism provides robustness to proportional hazards misspecification while retaining the efficiency and interpretability of classical models when their assumptions are approximately satisfied.

### 2.6. Iterative Survival–Censoring Ensemble Optimization

The IPCW loss functions introduced in Section 2.3 depend on accurate estimation of the censoring survival function *G*(*t* | ***X***). Rather than relying on a single parametric censoring model, **SuperSurv** optionally estimates the censoring distribution using a second Super Learner ensemble. This follows the joint survival–censoring stacking framework proposed by Westling *et al*. (2024), in which the survival distribution *S*(*t* | ***X***) and the censoring distribution *G*(*t* | ***X***) are estimated simultaneously through an iterative procedure.

The key idea is that estimation of *S*(*t* | ***X***) depends on the censoring model through IPCW weights, while estimation of *G*(*t* | ***X***) can similarly be expressed using pseudo-outcomes constructed from the current estimate of the survival distribution. Consequently, the two ensembles are updated alternately until convergence.

Specifically, let 𝒦_*S*_ and 𝒦_*G*_ denote candidate learner libraries for the survival and censoring distributions, respectively. Cross-validated predictions from these learners are first obtained on a common evaluation time grid 𝒯. The algorithm then iteratively updates the ensemble weights for *S*(*t* | ***X***) and *G*(*t* | ***X***) using pseudo-outcomes derived from the current estimates of the opposite distribution.

The ensemble weights are estimated by minimizing the cross-validated empirical loss subject to simplex constraints (nonnegative weights that sum to one). For the IPCW Brier score objective, this optimization reduces to a weighted least-squares problem, and the weights are estimated using nonnegative least squares (NNLS). Specifically, the cross-validated predictions are regressed on the IPCW pseudo-outcome under nonnegativity constraints, and the resulting coefficients are normalized to sum to one so that the final weight vector lies on the simplex 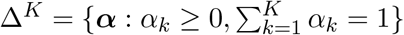.

For the IPCW negative log-likelihood objective, the loss function is no longer quadratic, and the ensemble weights are obtained by minimizing the cross-validated cross-entropy loss using constrained numerical optimization over the simplex.

Following Westling *et al*. (2024), the candidate learners themselves are fitted only once during the initial cross-validation stage. Subsequent iterations involve only the estimation of ensemble weights, making the procedure computationally efficient.

In addition to the standard ensemble estimator, **SuperSurv** also provides an optional model-selection strategy that returns the single best performing learner. Specifically, users may specify `selection = “best”`, which selects the candidate learner with the smallest cross-validated risk and assigns it weight 1, with all other weights set to zero. This corresponds to a hard model selection procedure, whereas the default `selection = “ensemble”` option estimates a convex combination of learners that typically achieves lower prediction error by averaging across models.

## 3. Software Design and Usage

The SuperSurv package provides a unified interface for survival ensemble learning built on the methodology described in Section 2. In addition to implementing the core Super Learner framework, the package includes utilities for model tuning, variable screening, performance evaluation, visualization, and model interpretation.

This section describes the overall package architecture and highlights several tools designed to facilitate practical survival modeling workflows.

### 3.1. Package Architecture

A major practical challenge in survival machine learning is the lack of a unified software interface. Popular algorithms such as Cox regression, random survival forests, gradient boosting, and penalized regression are implemented in different R packages, each requiring distinct data formats and producing predictions with incompatible output structures.

#### Algorithm 2

Iterative survival–censoring stacking

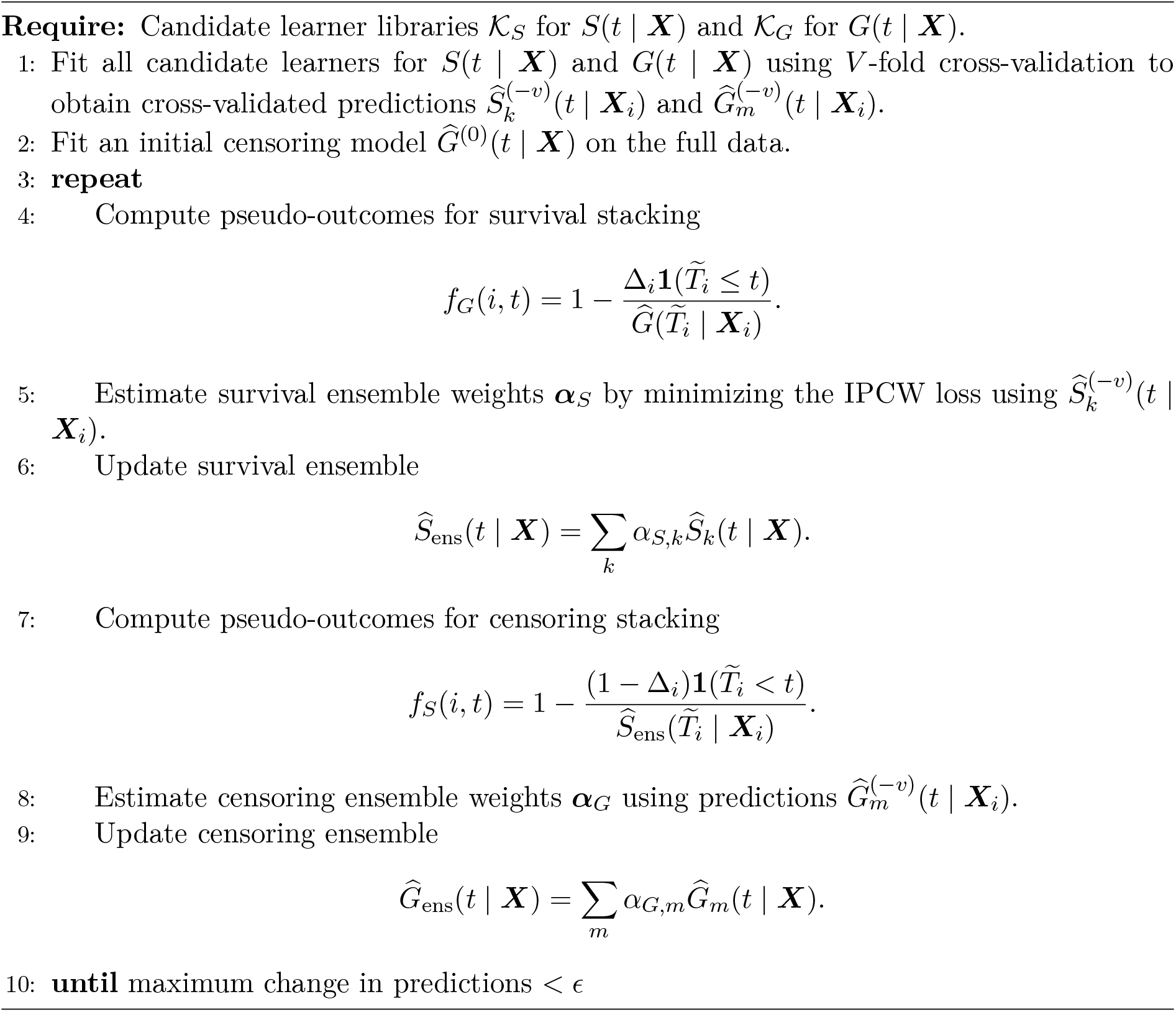

**SuperSurv** addresses this challenge by introducing a modular wrapper architecture that standardizes how survival learners are trained and how predictions are returned. Each base learner is implemented as a wrapper function with the prefix surv.*. These wrappers internally call external packages (e.g., **randomForestSRC, xgboost, glmnet**), but expose a unified interface to the **SuperSurv** framework.

The wrapper system ensures that all models return survival probability predictions on a common time grid. Regardless of the underlying algorithm, prediction outputs are converted into aligned matrices with dimension *N* ×|*𝒯*|, where *N* denotes the number of observations and 𝒯 denotes the evaluation time grid introduced in Section 2.3. This harmonized structure allows heterogeneous learners to be combined within the Super Learner ensemble.

Available learners and screening algorithms can be inspected directly within the package:

~~~
*R> # Install from GitHub
R> devtools::install_github(“yuelyu21/**SuperSurv**”)
R> library(“SuperSurv”)
R> library(“survival”)
R> list_wrappers()*
--- Prediction Models (surv.*) ---
 [1] “surv.aorsf” “surv.bart” “surv.coxph” “surv.glmnet”
 [5] “surv.parametric” “surv.rfsrc” “surv.weibull” “surv.xgboost” …
--- Screening Algorithms (screen.*) ---
[1] “screen.all” “screen.elasticnet” “screen.marg” “screen.var”
~~~

Prediction models are implemented as surv.* wrappers, while feature screening procedures follow the `screen`.* convention. This modular design allows new algorithms to be integrated into the framework by simply implementing a compatible wrapper that returns predictions in the required format.

Each wrapper follows a standardized interface that allows heterogeneous survival learners to be integrated seamlessly into the Super Learner framework. The design of this wrapper interface is described in Section 3.2.

### 3.2. Model Libraries and Wrapper Interface

All base learners in **SuperSurv** are implemented using a standardized wrapper interface. Each learner is defined as a function with the prefix `surv`.*. These functions serve as adapters that translate the input data into the format required by the underlying modeling package and convert model predictions into a standardized representation.

Specifically, each wrapper returns two components:

- `fit:` the fitted model object produced by the underlying algorithm.
- `pred:` a prediction matrix of dimension *N* × |𝒯 |, where *N* is the number of observations and 𝒯 is the evaluation time grid.

This contract ensures that predictions from heterogeneous algorithms can be directly stacked within the Super Learner meta-learning stage described in Section 2.3.

Prediction methods follow the standard R S3 convention using `predict.surv`.* methods, which accept a `newdata` argument and return survival probability predictions on the common time grid.

~~~
*R> # Fit a single wrapped base learner
R> fit0 <-surv.rfsrc(time = train$time,
+         event = train$event,
+         X = train[, covariate_cols],
+         newdata = test[, covariate_cols],
+         new.times = t_grid)
R> # Wrapper returns a standardized prediction matrix on the common grid
R> dim(fit0$pred)*
[1] n_test length(t_grid)
~~~

### 3.3. Hyperparameter Tuning and Grid Search

Many survival learning algorithms require tuning of hyperparameters that strongly influence predictive performance. To facilitate systematic hyperparameter exploration, **SuperSurv** provides a dynamic grid-generation utility that automatically constructs multiple wrapper functions corresponding to different parameter combinations.

The function `create_grid()` generates new learner wrappers by expanding a user-specified hyperparameter grid and programmatically creating wrapper functions with fixed parameter values. Each generated learner is assigned a descriptive name encoding its hyperparameter configuration.

~~~
*R> rf_tuning_params <-list(
+ nodesize = c(3, 15, 30),
+ ntree = c(100, 500)
+ )
R> rf_grid <-create_grid(
+ base_learner = “surv.rfsrc”,
+ grid_params = rf_tuning_params
+ )
R> rf_grid*
[1] “surv.rfsrc_nodesize3_ntree100”
[2] “surv.rfsrc_nodesize15_ntree100”
[3] “surv.rfsrc_nodesize30_ntree100”
[4] “surv.rfsrc_nodesize3_ntree500”
[5] “surv.rfsrc_nodesize15_ntree500”
[6] “surv.rfsrc_nodesize30_ntree500”
~~~

These dynamically generated learners can then be included directly in the Super Learner library. Because each configuration is treated as an independent base learner, the meta-learning stage automatically identifies the optimal hyperparameter setting through cross-validated risk minimization.

### 3.4. Variable Screening

High-dimensional survival prediction often involves settings where the number of covariates *p* is large relative to the sample size *n* (e.g., molecular profiles or large EHR feature sets). To reduce computational burden and mitigate overfitting, **SuperSurv** supports an optional screening stage that selects a subset of covariates prior to fitting each base learner. This design follows the screening-then-estimation paradigm used in survival Super Learner implementations, where different learners may employ different screening rules.

Let ***X*** ∈ ℝ^n×p^ denote the training covariate matrix. A screening algorithm produces a binary inclusion vector

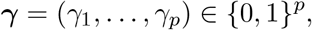

where *γ*_j_ = 1 indicates that covariate *j* is retained. The screened design matrix is then ***X***_γ_, consisting of the subset of columns with *γ*_j_ = 1. In **SuperSurv**, screening is performed *within each cross-validation training fold* to avoid information leakage.

#### Implemented screeners

In addition to the marginal Cox screening and penalized Cox (LASSO/Elastic Net) screening commonly used in survival Super Learner libraries, **SuperSurv** extends the screening library with both supervised and unsupervised alternatives: (i) `screen.all`, which retains all covariates; (ii) `screen.marg`, which ranks features by univariate Cox association; (iii) `screen.glmnet` and `screen.elasticnet`, which fit penalized Cox models and retain covariates with nonzero coefficients; (iv) `screen.rfsrc`, which fits a fast random survival forest and retains covariates with positive variable importance (VIMP); and (v) `screen.var`, which retains the top fraction of covariates by empirical variance, providing an unsupervised filter suitable for noisy high-dimensional inputs.

To ensure stability, each screener enforces a minimum number of retained variables (argument `minscreen`), so that the downstream learner is always estimable even when all features appear weak in finite samples.

#### Screened learner libraries

A screened library is specified as a list of learner–screener pairs, allowing different learners to use different screening rules:

~~~
*R> my_screen_library <-list(
+   c(“surv.coxph”,   “screen.all”),
+   c(“surv.coxph”,   “screen.marg”),
+   c(“surv.weibull”, “screen.elasticnet”),
+   c(“surv.rfsrc”,   “screen.var”)
+ )*
~~~

The selected covariates within each fold are stored (optionally) to enable reproducibility and post hoc inspection of the screening behavior.

### 3.5. Model Evaluation Metrics

**SuperSurv** provides a unified set of evaluation utilities for comparing the fitted ensemble against each candidate base learner on a common time grid 𝒯 = {*t*_1_, …, *t*_m_}. Given test data 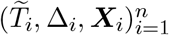 and predicted survival probabilities 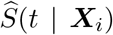 evaluated at *t* ∈ 𝒯, the package computes time-dependent performance curves based on three widely used metrics: the IPCW Brier score, cumulative/dynamic time-dependent AUC, and Uno’s C-index.

#### Time-dependent IPCW Brier score

For each evaluation time *t* ∈ *𝒯*, the IPCW Brier score is computed as an empirical risk over the expanded (*i, t*) set:

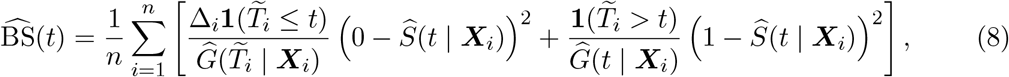

where 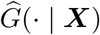 denotes an estimator of the censoring survival function. Lower values indicate better accuracy of the predicted survival probabilities.

#### Time-dependent AUC

To assess time-varying discrimination, **SuperSurv** computes a cumulative/dynamic AUC curve across based 𝒯 on time-dependent ROC methodology for right-censored outcomes. Conceptually, the AUC at time *t* measures the ability of the model’s risk ranking (equivalently, low 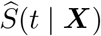 ) to separate individuals who experience the event by time *t* from those who remain event-free beyond *t*, while accounting for censoring.

#### Uno’s C-index

As a complementary measure of concordance that remains well-defined under heavy censoring, **SuperSurv** provides Uno’s C-index evaluated at each time point *t* ∈ *𝒯*, producing a longitudinal concordance curve. Higher values indicate better discrimination.

#### Benchmarking the ensemble against base learners

The primary benchmarking utility, `plot_benchmark()`, constructs longitudinal performance curves for the ensemble and each base learner using predictions returned by `predict()` evaluated on 𝒯. This comparison is implemented in a fully model-agnostic manner: any fitted object that supports `predict(object, newdata, new.times)` and returns an *n* × *m* survival probability matrix can be benchmarked under the same interface.

~~~
*R> plot_benchmark(
+ object = fit_supersurv,
+ newdata = X_te,
+ time = test$duration,
+ event = test$event,
+ eval_times = new.times)*
~~~

#### Visualization of individual survival curves

To facilitate clinical interpretation at the individual level, `plot_predict()` visualizes predicted survival trajectories 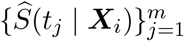 for one or more patients using step-function plots consistent with the discrete evaluation grid.

~~~
*R> preds <-predict(fit_supersurv, newdata = X_te, new.times = new.times)
R> plot_predict(preds = preds, eval_times = new.times, patient_idx = 1:6)*
~~~

#### Calibration assessment at a fixed time horizon

In addition to discrimination and overall accuracy, **SuperSurv** includes a pragmatic calibration diagnostic `via plot_calibration()` at a user-specified horizon *t*^⋆^. The function partitions individuals into quantile bins of predicted survival 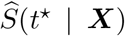, then compares the mean predicted survival in each bin to the observed Kaplan–Meier estimate at *t*^⋆^ within that bin. The resulting reliability curve is plotted against the 45^°^ line representing perfect calibration.

~~~
*R> plot_calibration(
+ object = fit_supersurv,
+ newdata = X_te,
+ time = test$duration,
+ event = test$event,
+ eval_time = 150,
+ bins = 5)*
~~~

### 3.6. Model Interpretation and Explainability

To support transparent reporting and clinical interpretability, **SuperSurv** provides a unified explainability layer for both ensemble models and individual base learners. The primary goal is to explain a *scalar risk functional* derived from a fitted survival model, denoted 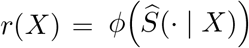, where *ϕ*(·) may represent, for example, risk at a prespecified horizon *t*^⋆^ or another user-defined summary of the predicted survival curve. This design allows model-agnostic explanations even when base learners differ in their internal parameterization.

#### Kernel SHAP for global and local explanations

The function `explain_kernel()` interfaces with the fastshap package to compute Kernel SHAP values for *r*(*X*) using Monte Carlo Shapley estimation with a user-specified background dataset. For a fitted model *r*(·) and feature set {1, …, *p*}, SHAP decomposes the prediction into additive feature contributions

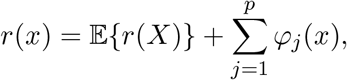

where *φ*_j_(*x*) is the Shapley value for feature *j* at observation *x*.

#### Weighted SHAP for ensembles

For a convex Super Learner ensemble, **SuperSurv** optionally computes SHAP values for each *active* base learner and aggregates them using the estimated meta-learner weights. Specifically, if the explained risk functional is linear in the library predictions,

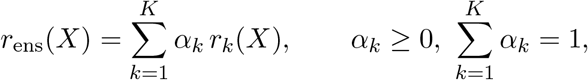

then the linearity property of Shapley values implies 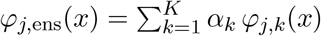 . In practice, **SuperSurv** computes Kernel SHAP for each base learner *k* and returns the weight-averaged SHAP matrix, enabling global importance ranking (mean |*φ*_j_| ), cohort-level summary plots (beeswarm), and patient-level explanations (waterfall plots).

~~~
*R> shap_vals <-explain_kernel(
+ model = fit_sl,
+ X_explain = X_test[1:50, ],
+ X_background = X_train[1:100, ],
+ nsim = 20)*
~~~

#### Time-dependent explainability via survex

Because survival prediction is intrinsically time-indexed, **SuperSurv** also provides `explain_survex()`, which constructs a standard survex explainer object by supplying a unified survival-prediction wrapper. This bridge unlocks the full survex ecosystem for time-dependent diagnostics, including dynamic permutation importance, time-indexed partial dependence profiles, and local SurvSHAP(*t*) explanations of how covariates shift the predicted survival trajectory across a prespecified grid of times.

~~~
*R> surv_explainer <-explain_survex(
+ fit_sl,
+ data = X_test,
+ y = Surv(time, event),
+ times = grid) R> library(survex)
R> plot(model_parts(surv_explainer))*
~~~

### 3.7. Adjusted Marginal Contrasts via Restricted Mean Survival Time (RMST)

Although hazard ratios from proportional hazards models remain a common reporting standard, they can be difficult to interpret on an absolute time scale and may be unstable under non-proportional or crossing hazards. To provide an estimand on the survival-time scale, **SuperSurv** implements restricted mean survival time (RMST)(Royston and Parmar 2013) contrasts using g-computation (standardization) on top of the fitted Super Learner survival model.

For a prespecified horizon *τ*, the individual-level RMST is defined as the area under the survival curve,

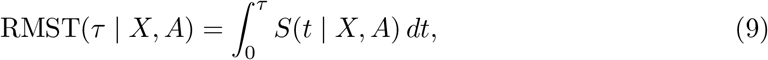

which **SuperSurv** evaluates numerically on the prediction grid using a left Riemann sum.

Given a binary grouping variable or exposure *A* ∈ {0, 1} included among the covariates, **SuperSurv** estimates a standardized marginal contrast on the RMST scale by comparing the predicted restricted mean survival under the two levels of *A*. Specifically, the target contrast is

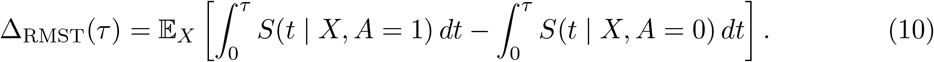

#### Standardized RMST contrast

Let *A* ∈ {0, 1} denote a binary grouping variable included among the model covariates. **SuperSurv** estimates the standardized mean RMST under the two levels of *A* by (i) setting *A* = 1 for all individuals, predicting 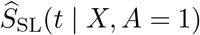, and integrating to obtain 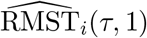 ; and (ii) repeating with *A* = 0 to obtain 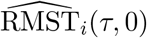 . The resulting marginal contrast is

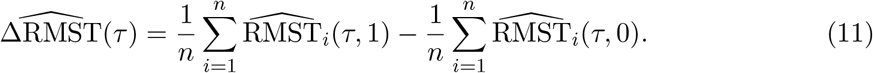

Equivalently, defining the individual-level predicted contrast

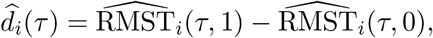

the plug-in estimator can be written as

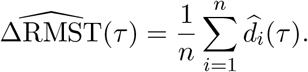

This quantity should generally be interpreted as an adjusted marginal contrast on the RMST scale. When *A* corresponds to a manipulable intervention and the necessary identification assumptions hold, the same standardized procedure may also support a causal interpretation. For non-manipulable attributes such as sex or biomarker status, however, 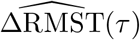 should be interpreted as a covariate-adjusted group contrast rather than a causal effect.

#### Perturbation-based inference

To provide uncertainty quantification for the standardized RMST contrast, **SuperSurv** optionally implements a perturbation-based inference procedure conditional on the fitted ensemble model(Yang 2013; Jin, Ying, and Wei 2001). For perturbation replicate *b* = 1, …, *B*, generate independent positive random weights

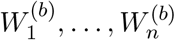

with mean 1 (e.g., exponential weights), and apply these weights to the *n* individuals respectively to do a weighted data analysis in a similar fashion as described above to obtain *d*_i_(*τ* )^(b)^, *i* = 1,, *n*, then define the perturbed RMST contrast

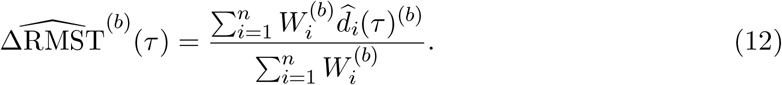

Let

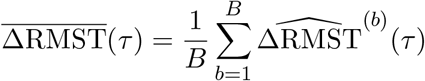

denote the average perturbed contrast. The perturbation-based standard error is estimated by

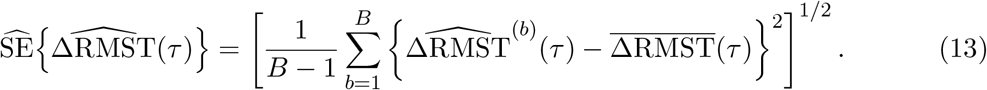

A Wald-type 100(1 − *α*)% confidence interval is then given by

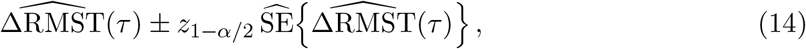

and the corresponding two-sided Wald statistic for testing

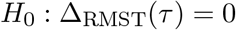

is

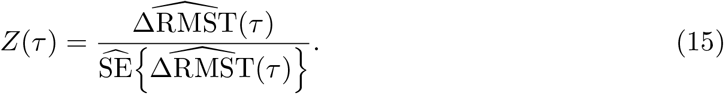

The p value is computed as

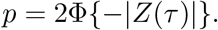

In the current implementation, this perturbation procedure holds the fitted **SuperSurv** model fixed, including the selected learner library, hyperparameters, fitted base learners, and ensemble weights. Thus, the resulting uncertainty quantification is conditional on the final fitted ensemble and reflects variability in the standardized RMST contrast given the estimated survival model.

~~~
*R> # Adjusted RMST contrast with perturbation-based inference
R> rmst_res <-estimate_marginal_rmst(
+ fit = fit_sl,
+ data = clinical_data,
+ trt_col = “treatment_status”,
+ times = prediction_grid,
+ tau = 100,
+ inference = TRUE,
+ B = 200,
+ seed = 123
+ )*
~~~

#### Effect curves and diagnostics

To assess how the adjusted contrast evolves with the restriction time, **SuperSurv** provides `plot_marginal_rmst_curve()`, which evaluates 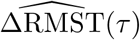 over a user-specified grid of *τ* values. For a simple model-checking diagnostic, `plot_rmst_vs_obs()` compares each individual’s predicted RMST against their observed follow-up time, with visual stratification by event status.

## 4. Empirical Application: METABRIC Breast Cancer Data

To demonstrate an end-to-end survival prediction workflow, we apply **SuperSurv** to the METABRIC breast cancer dataset included with the package. This example highlights three core strengths of the framework: (i) a unified API for training heterogeneous survival learners under right censoring, (ii) automated ensemble construction with time-dependent benchmarking, and (iii) downstream interpretability and clinically meaningful marginal contrasts using RMST.

### 4.1. Data Preparation and Reproducible Initialization

We begin with a standard 70%*/*30% train–test split. Predictors are selected as the covariate block indexed by the prefix x, and evaluation times are defined on a clinically interpretable grid.

~~~
*R> library(SuperSurv)
R> library(survival)
R> data(“metabric”, package = “SuperSurv”)
R> set.seed(42)
R> train_idx <-sample(seq_len(nrow(metabric)), size = 0.7 * nrow(metabric))
R> train <-metabric[train_idx, ]
R> test <-metabric[-train_idx, ]
R> x_cols <-grep(“^x”, names(metabric), value = TRUE)
R> X_tr <-train[, x_cols]
R> X_te <-test[, x_cols]
R> new.times <-seq(30, 300, by = 30)*
~~~

### 4.2. Library Construction and Hyperparameter Tuning

We construct an event-model library that spans interpretable parametric and semiparametric baselines as well as a flexible nonparametric learner. In particular, we include (i) a Cox model (surv.coxph), (ii) a Weibull model (surv.weibull), and (iii) a tuned family of Random Survival Forests generated automatically via create_grid(). For the censoring model used for IPCW, we use a Cox model for stability.

~~~
*R> rf_grid <-create_grid(
+ base_learner = “surv.rfsrc”,
+ grid_params = list(nodesize = c(5, 20), ntree = 100)
+ )
R> event_lib <-c(“surv.coxph”, “surv.weibull”, rf_grid)
R> cens_lib <-c(“surv.coxph”)*
~~~

### 4.3. Ensemble Training and Weight Inspection

We fit the meta-learner using *K*-fold cross-validation. To enable downstream interpretability and RMST-based contrasts that require refitting/prediction under counterfactual covariate settings, we save the fitted base-learner library via control = list(saveFitLibrary = TRUE). If computational resources permit, cross-validation can be parallelized using future; here we illustrate a lightweight configuration.

~~~
*R> fit_sl <-SuperSurv(
+ time = train$duration,
+ event = train$event,
+ X = X_tr,
+ newdata = X_te,
+ new.times = new.times,
+ event.library = event_lib,
+ cens.library = cens_lib,
+ selection = “ensemble”,
+ control = list(saveFitLibrary = TRUE),
+ nFolds = 5,
+ verbose = FALSE
+ )
R> round(fit_sl$event.coef, 3)*
surv.coxph_screen.all                           surv.weibull_screen.all
                                         0.587                                   0.000
surv.rfsrc_nodesize5_ntree100_screen.all        surv.rfsrc_nodesize20_ntree100_screen.all
                                         0.068                                   0.345
~~~

The resulting weight vector provides a transparent summary of which model classes contribute most to out-of-sample predictive performance in this dataset.

### 4.4. Time-Dependent Predictive Benchmarking

We evaluate the fitted ensemble on the held-out test set using the package’s built-in bench-marking utilities, which compute time-dependent prediction metrics (e.g., IPCW Brier score, time-dependent AUC, and Uno’s C-index) on a common time grid.

~~~
*R> plot_benchmark(
+ object = fit_sl,
+ newdata = X_te,
+ time = test$duration,
+ event = test$event,
+ eval_times = new.times
+ )*
~~~

**Figure 2:**
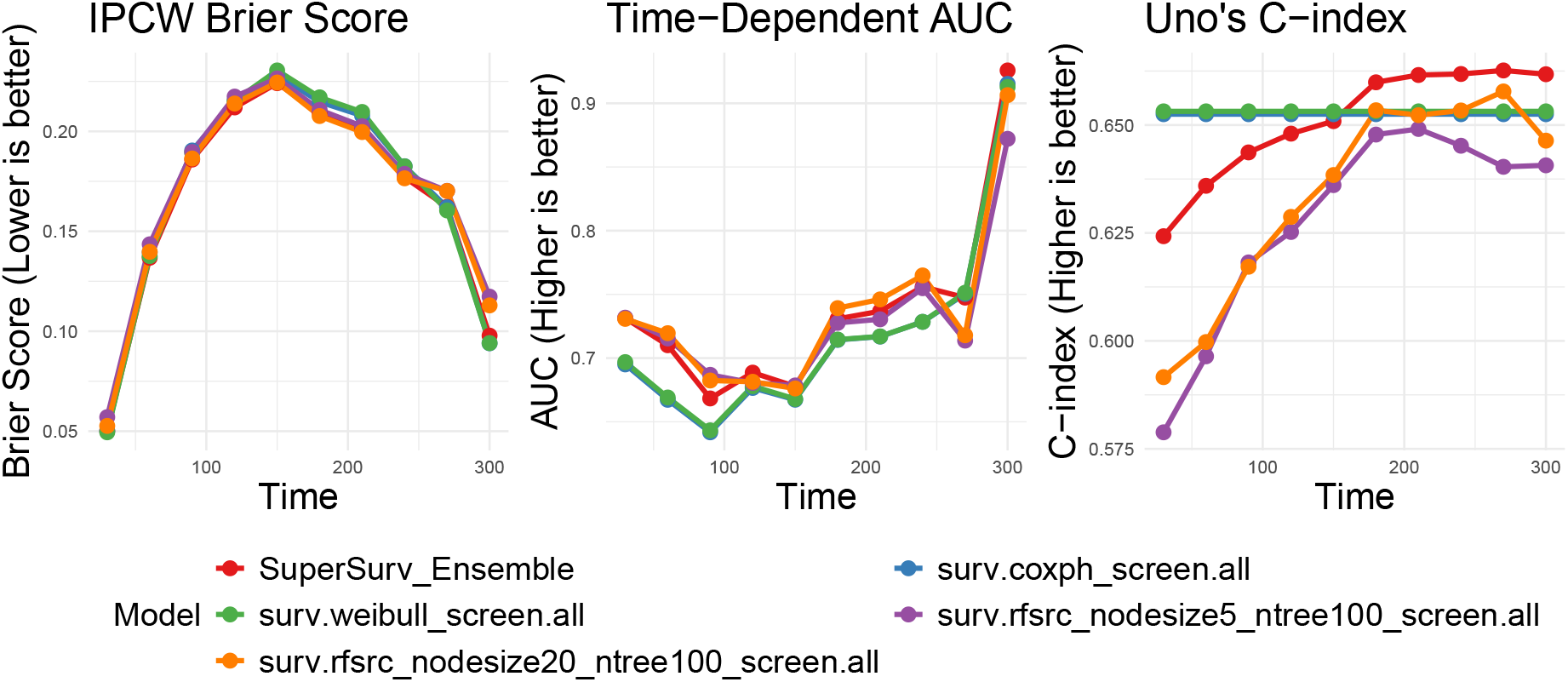
Time-dependent benchmarking on the METABRIC test set comparing the SuperSurv ensemble with each base learner.

### 4.5. Explainable AI via Kernel SHAP

To provide model transparency, we compute Kernel SHAP values for a subset of test patients using a subset of training patients as a reference distribution. Because **SuperSurv** forms an ensemble of base learners, SHAP values can be computed at the learner level and aggregated according to the fitted ensemble weights.

~~~
*R> shap_vals <-explain_kernel(
+ model = fit_sl,
+ X_explain = X_te[1:50, ],
+ X_background = X_tr[1:100, ],
+ nsim = 10
+ )
R> plot_beeswarm(shap_vals, data = X_te[1:50, ], top_n = 8)*
~~~

**Figure 3:**
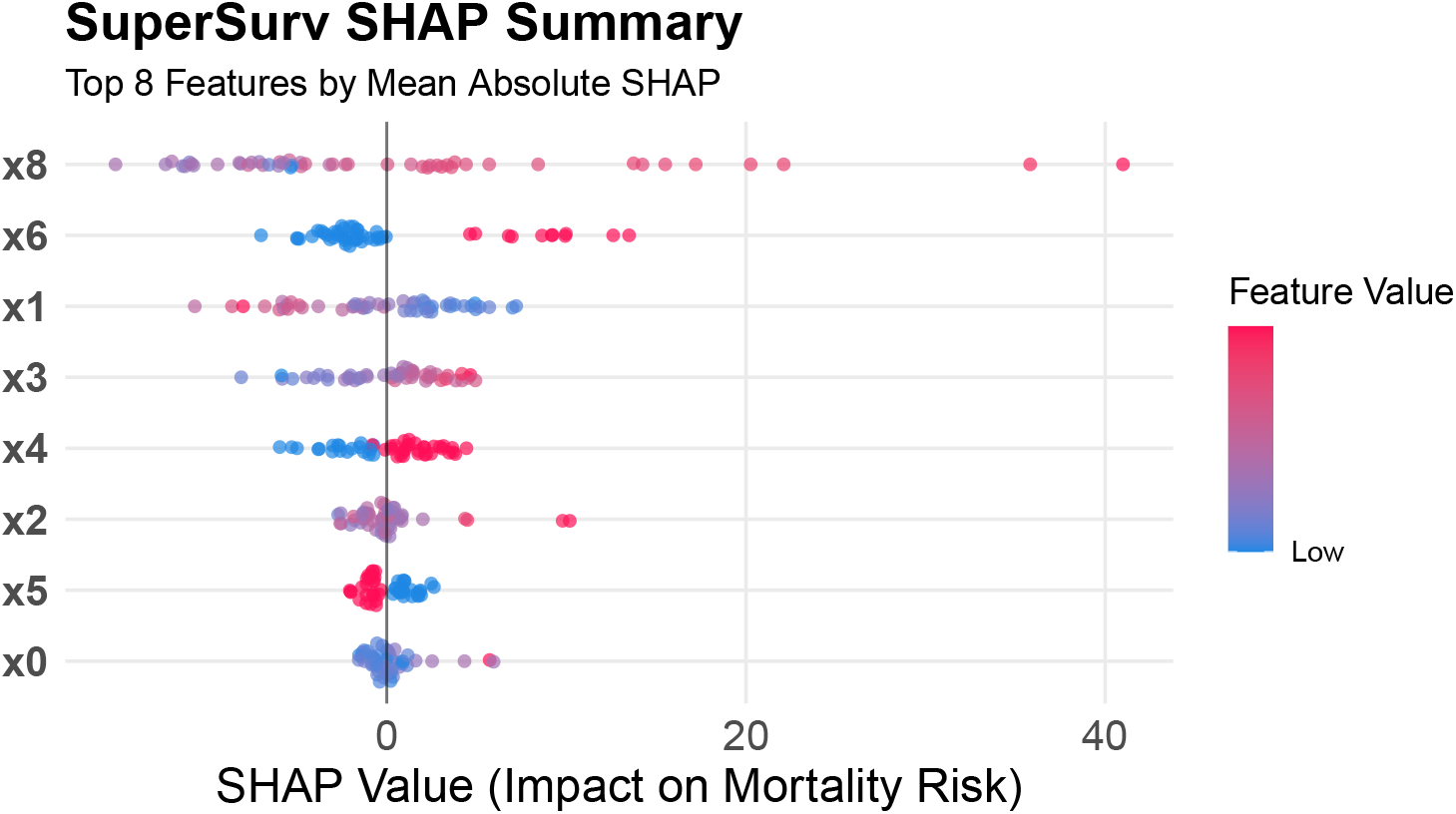
SHAP beeswarm plot for the fitted **SuperSurv** ensemble showing the direction and magnitude of the top contributing features to mortality risk.

### 4.6 Covariate-Adjusted Marginal Contrast via RMST

Finally, we illustrate how the fitted ensemble can be used to report an absolute, clinically interpretable contrast on the survival-time scale using restricted mean survival time (RMST). Treating `x4` as a binary exposure (e.g., a biomarker), we estimate the covariate-adjusted marginal contrast in RMST up to *τ* = 100.

~~~
*R> rmst_results <-estimate_marginal_rmst(
+ fit = fit_sl,
+ data = metabric,
+ trt_col = “x4”,
+ times = new.times,
+ tau = 100,
+ inference = TRUE,
+ B = 200,
+ seed = 123
+ )
R> rmst_results$ATE_RMST
R> rmst_results$SE_RMST
R> rmst_results$CI_RMST
R> format.pval(rmst_results$p_value, digits = 3, eps = 1e-16)*
[1] -1.744314
[1] 0.02713712
    lower      upper
-1.797502 -1.691126
[1] “< 1e-16”
*R> > p1 <-plot_marginal_rmst_curve(
+ fit = fit_sl,
+ data = metabric,
+ trt_col = “x4”,
+ times = new.times,
+ tau_seq = seq(50, 300, by = 50),
+ inference = TRUE,
+ B = 200,
+ seed = 123,
+ ci_level = 0.95
+ )
R> p2 <-plot_rmst_vs_obs(
+ fit = fit_sl,
+ data = metabric,
+ time_col = “duration”,
+ event_col = “event”,
+ times = new.times,
+ tau = 350
+ )
R> p1 + p2*
~~~

**Figure 4:**
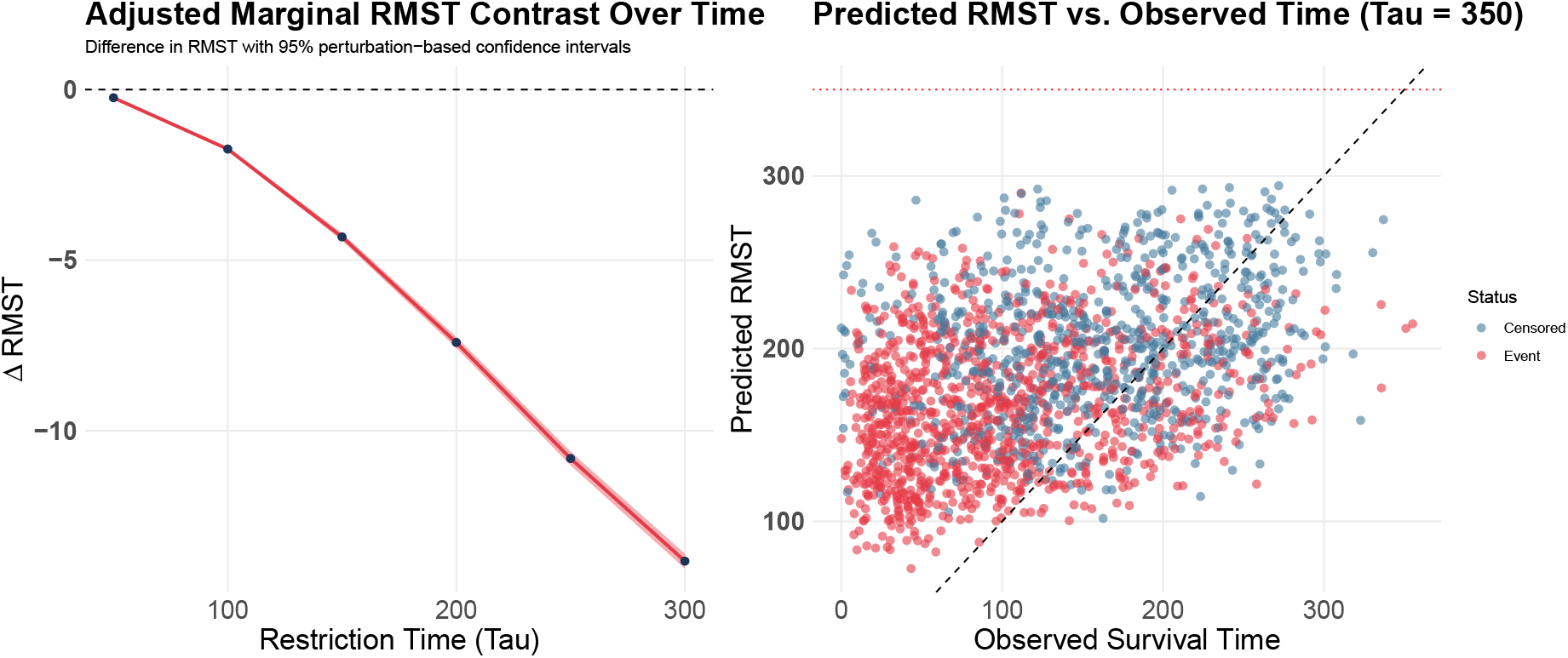
RMST-based post hoc summaries from the fitted **SuperSurv** ensemble. Left: adjusted marginal RMST contrast across restriction times, with optional perturbation-based confidence intervals. Right: predicted RMST versus observed follow-up time, shown as a simple diagnostic of agreement between restricted survival predictions and observed outcomes.

This RMST-based contrast summarizes the group difference on an absolute time scale and can remain meaningful under non-proportional hazards patterns where hazard ratios may be difficult to interpret.

Together, these steps demonstrate how **SuperSurv** enables a complete survival modeling workflow—from ensemble construction and performance benchmarking to explainable AI and clinically interpretable survival contrasts—within a unified software framework.

## 5. Discussion

This article introduced **SuperSurv**, an R package for ensemble-based survival modeling built on the Super Learner framework. The package provides a unified infrastructure for combining heterogeneous survival algorithms, performing cross-validated risk minimization, and producing time-dependent survival predictions within a standardized API.

A central contribution of **SuperSurv** is the integration of diverse survival model classes—including classical statistical models, tree-based learners, and modern machine learning algorithms— within a unified ensemble architecture. By estimating convex combinations of candidate learners, the Super Learner ensemble adapts to the underlying structure of the data and can achieve strong predictive performance across a wide range of survival modeling scenarios.

The framework is flexible and allows users to incorporate automated hyperparameter tuning, feature screening algorithms, and alternative loss functions such as IPCW log-loss.

Beyond prediction accuracy, **SuperSurv** emphasizes interpretability and practical usability. The package provides built-in tools for explainable machine learning, including Kernel SHAP and integration with the survex ecosystem for time-dependent explanation methods such as SurvSHAP(*t*). These tools enable users to investigate both global feature importance and patient-level explanations of survival predictions.

In addition, **SuperSurv** supports downstream survival contrasts based on restricted mean survival time (RMST). By combining Super Learner survival predictions with standardization (g-computation), the package allows researchers to estimate covariate-adjusted differences in expected survival time between groups. This approach provides an interpretable alternative to hazard ratios and remains valid even when the proportional hazards assumption does not hold.

The empirical application using the METABRIC breast cancer dataset illustrates how SuperSurv can be used to construct a complete survival analysis workflow, including ensemble construction, predictive benchmarking, model interpretation, and estimation of adjusted survival contrasts. This end-to-end example highlights the package’s ability to bridge predictive modeling and clinically interpretable inference.

Several directions remain for future development. Potential extensions include support for competing risks models, multi-state survival processes, and causal inference frameworks based on targeted learning. In addition, further integration with distributed computing frameworks could enable scalable training for extremely large biomedical datasets.

Overall, **SuperSurv** provides a flexible and extensible toolkit for modern survival analysis, combining the statistical guarantees of Super Learner with practical tools for prediction, interpretation, and clinically meaningful survival contrasts.

